# Habituation and goal-directed arbitration mechanisms and failures under partial observability

**DOI:** 10.1101/2020.11.24.396630

**Authors:** Martí Sánchez-Fibla

## Abstract

We often need to make decisions under incomplete information (partial observability) and the brain manages to add the right minimal context to the decision-making. Partial observability may also be handled by other mechanisms than adding contextual experience / memory. We propose that parallel and sequential arbitration of Habituation (Model-Free, MF) and Goal-Directed (Model-Based, MB) behavior may be at play to deal with partial observability “on-the-fly”, and that MB may be of different types (going beyond the MF/MB dichotomy [4]). To illustrate this, we identify, describe and model with Reinforcement Learning (RL) a behavioral anomaly (an habituation failure) occurring during the so-called Hotel Elevators Rows (HER, for short) task: a prototypical partial observation situation that can be reduced to the well studied Two and One Sequence Choice Tasks. The following hypothesis are supported by RL simulation results: (1) a parallel (semi)model-based successor representation mechanism is operative while learning to habituate which detects model-based mismatches and serves as an habituation surveillance, (2) a retrospective inference is triggered to identify the source of the habituation failure (3) a model-free mechanism can trigger model-based mechanisms in states in which habituation failed. The “failures” in the title refer to: the habituation failures that need to be monitored and surveilled (1) and to the failures that we identified in prototypical state of the art Model-Based algorithms (like DynaQ) when facing partial observability. As other research on MF/MB arbitration shows, the identification of these new mechanisms could shine light into new treatments for addiction, compulsive behavior (like compulsive checking) and understand better accidents caused by habituation behaviors.

## 1 Introduction

The underlying neuro-physiological mechanisms of how the brain interleaves and jointly controls habitual and goal-directed behavior [5] are just starting to be unraveled [9] and modeled [6] with Reinforcement Learning (RL) [18, 21, 27, 30]. These two mechanisms of behavioral control were first related to Model-Free (MF) and Model-Based (MB) RL in [7]^3^ and beyond RL, have been associated with Kahneman’s Fast System 1, Slow System 2 [17], Learners versus Solvers [14], automatic/stimulus-driven responses versus deliberative reasoning, etc. Regarding RL as a modeling framework of cognitive processes, the MF/MB dichotomy may be too simplistic [4], needing to accommodate inbetween approaches like Successor Representations [24, 26, 28], various typologies of MB processes like retrospective inference [27] and a richer landscape of MF/MB arbitration. Daw et al. [6] characterized with a human study the MF and MB contributions used in behavioral control via a Sequential Two-choice task. Their study reveals that people enforce both MF and MB strategies at the same time in parallel (as found also in [30]). As suggested by [11] the direct relationship between Habitual and Model-Free may be even too simplistic as Goal-Directed behavior may become automatic as well.

It is widely assumed^4^ that the brain uses Goal-Directed/MB strategies in novel environments/tasks and tries to enforce Habitual/MF-RL when exposed to repetitions of the same task characteristics, with the aim of freeing resources (being MB strategies more costly). Moreover, when habits have been created (for example in usual daily travels) they are difficult to suppress [1] and the effect is stronger with cognitive load and with the presence of time constraints [16]. The brain operates in the regime of optimizing the speed-accuracy trade-off [18] of Habitual and Goal-Directed behaviors. It acts as an habituation machine, that is constantly trying to off-load tasks from being “consciously monitored”. But, this habituation process can fail abruptly (like we will argue, in the case of the task and anomaly that we present), or fail constantly/smoothly in the case of the Sequential Two-Choice Task [6], as a dynamic mechanism for reward switching is imposed, which turns the environment to be dynamic and non-stationary. Since [6] introduced the Two-Choice Task behavioral paradigm to measure the amount of Habitual/MF, Goal-Directed/MB employed by participants, its adequacy has been debated [2, 19].

In this paper we present and model computationally (based on RL) an observed human behavioral anomaly that we call *Hotel Elevator’s Rows* (HER for short) task and anomaly (see Figure **1**), in which due to partial observability (incomplete task/sensory information), the brain can fail repeatedly to apply an acquired habitual behavior. Goal-Directed control must come into rescue while building an “alarm” detector for anticipating the potential future failures of the automatic control. These alarm warnings can be characterized as MF associations with a particular state, in such a way that, when that state is encountered again, the alarm is raised and control is taken by a Goal Directed process with an increased level of attention. So in this case the arbitration between Habituation and Goal-Directed behavior can be produced with one shot learning, after an habituation failure. By failure we mean that an undesired state is reached, detected through a Model-Based mismatch based on Successor Representations (SRs). We show how, under partial observability this additional Model-Based expectancy signal (i.e. coming from SRs Temporal Difference learning) can be fundamental to stabilize Model-Free Q-Learning convergence. We highlight here the fact that is “one-shot” as the VPI measure (Value of Perfect Information introduced in [18]) being linked to uncertainty and variance in the learning may need several trials to build up and arbitrate between MF and MB systems. We also show that a prototypical Model Based algorithm like Prioritized Sweeping DynaQ (used in [21] for providing plausible models of various brain phenomena: like hippocampal forward sweeps / planning and replay) worsens its converge under the partial observability imposed by the HER task.

**Figure 1:**
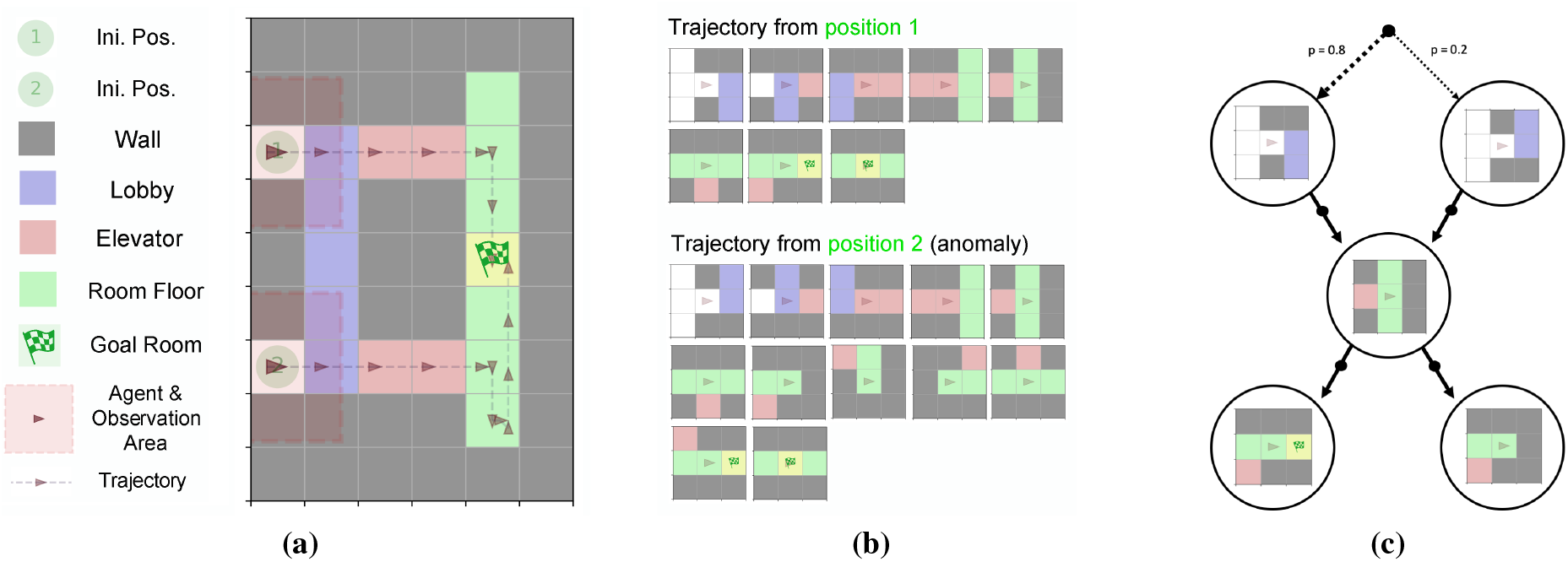
*Hotel Elevators Rows* task (HER for short). **(a)** The HER task as a gridworld. A 9×6 grid represents the environment in which the agent is depicted as a dark red triangle. Its observation area is depicted with a red shaded square around him. **(b)** Observations of the agent during two trajectories shown in (a). When the agents starts at position two (at the bottom) the so-called HER anomaly is experienced. The agent observation is delivered rotated when the agent turns; so the agent is centered in the observation always facing right. **(c)** One-Choice version of the HER task described in Section 3.1.

Its only recently that the arbitration of MF and MB mechanisms in the brain, have been studied under incomplete information [27] (also referred as partial observability). For instance, behavioral research derived from [6] only considers perfect information tasks [2, 19]. Computational modeling research [21, 28] has also only considered global states, or complete information states in their RL models and formulations. Partial observability (or incomplete information) is a very common situation in decision making from a behavioral perspective; but from a computational point of view as well, as current Deep Neural Network approaches, make approximations of the state space that induce an uncontrolled partial observability. A recent computational approach [33] explores another side of partial observability, noisy uncertain observations and not incomplete information, as we deal here). Non-stationarity, on the other hand, has received more attention. Adaptability to changing reward conditions as in [28], and the long list of papers that followed the Two-Choice task [6], all maintain a non-stationary changing reward to penalize habituation. The task that we model here is stationary and yet presents habituation and MB failures due to partial observability and is challenging in terms of MF/MB arbitration. We do not present behavioral results (only simulations), but we do propose several ideas on how to test the proposed principles with behavioral experiments.

A recent Model-Based RL survey [23] including a section of methods dealing with partial-observability only mentions memory-based RL (state history recording, recurrency and Neural Turing Machine approaches). Thus we consider a new direction as partial observability could also be addressed with on-the-fly behavioral mechanisms: a) habituation semi Model-Based monitoring (driven for example by Successor Representations mismatch detectors). b) retrospective Model-Based alarm reinforcement (adding up to other recently described retrospective mechanisms [27]) and c) Model Free alarm triggers (leading to increased attentional demands and planning mechanisms).

In summary, we propose and model the so-called HER partial observable task (introduced in next Section 1.1) that captures a recognizable human behavioral conflict between habitual and goal-directed behavior. In the process of learning to habituate the brain can fall in a partial observable trap. To capture all aspects of the equivalent human behavioral task we present four different HER environments (see Section 2.2) with: (1) global, complete information states, (2) incomplete information / partial observable states, (3) memory of last *n* experienced states, (4) complete/incomplete observation size on-demand. The HER anomaly is reproduced with all RL algorithms (MF Q-Learning, MB DynaQ and Reinforce Policy Gradient, see Section 3.2) in the partial observable (2) and Observation size on-demand (4) environments. We show that a prototypical Model-Based algorithm, Prioritized Sweeping DynaQ, which is the best performing method with complete/global information, fails to converge under partial observability (Section 3.6). We propose previously unreported arbitration mechanisms in the process of learning to habituate: Inline with [4], we present evidence for MF and MB parallel processes (as in [30], in our case, learning to habituate while being monitored by semi Model-Based state expectancy visitations). A Model-Based SR mechanism that acts as a mismatch detector and could operate at several dimensions: states expected to be visited and also in the correct timing (Section 3.8). We characterize Model-Based processes in different forms: sub-goal task division, increased attentional demands for enhanced state information, retrospective inference (as in [27]) for explaining the source of habituation failures (see Section 3.9) and reconsidering decisions and forward planning (as in [18]). Finally, we describe an MF mechanism that can remember states identified as sources of habituation failure for switching to Goal-Directed MB behavior when encountered again in the future (see Section 3.10). Altogether provide plausible solution of how we solve the HER anomaly in few shot exposures to an habituation failure.

We start with some preliminary high level descriptions of the HER task (next Section 1.1) and its different modeling paradigms that we will employ (Section 1.2).

### 1.1 The *Hotel Elevator’s Rows* (HER) task and anomaly

Lets describe a scenario in which Habitual and Goal-Directed behavior interact under incomplete sensory state information (partial observability). We’ll refer to it as the *Hotel Elevator’s Rows*. Imagine your are in the lobby of a hotel with two sides of elevators and the goal to get to your room. You have been a few days in the hotel and you already build a sort of habitual behavior and you tend to unconsciously prefer elevators that are on the left side. In that particular moment an elevator on the right side opens empty inviting you to a quicker ride to your room. After getting out of the elevator and executing your habitual behavior program, you find your self in the wrong corridor. What has happened here? And how can we model this anomaly?

When taking the uncommon elevator’ side, the habitual system is held back because you prioritize time and you want to get to the room as quick as possible. While being in the elevator your real position with respect to you room is not observable considering only the available visual information, thus we call it partially observable. When you exit the elevator, habitual behavior is executed and that brings you to the wrong corridor.

The executive brain has two possibilities to solve this situation if it repeats and you wish to avoid it. (a) Trying to remember from which side you took the elevator (1 bit of information) and sustain the context info during the elevator’s trip. (b) Being more careful when getting out of the elevator, anticipating the anomaly by searching for distal visual cues that will help distinguishing the correct corridor earlier on, thus grabbing your full attention and preventing you of other thoughts. For doing (b) it has to remember that there is something to be done when exiting the elevator due to previous repetitions of the anomaly.

Option (a) is a typical approach in Reinforcement Learning, any partial observable problem (containing states with incomplete information) can be solved with sufficient memory (see for example the mentioned review [23]). Option (b) plants the seed for a new approach to problems with incomplete information and argues for an “on-the-fly” mechanism to solve potential identified ambiguous and undesired situations which are the subject of the current paper.

### 1.2 HER task: high level modeling

We check-in to a new hotel and we have a certain a priori knowledge of how to subdivide the goal of getting to our assigned room into subgoals: get into to hall, get into the elevator go to the corresponding floor, turn left or right after exiting the elevator exit, navigate the right corridors until we get into our room number. This navigation is Goal-Directed and requires maximum attention, for instance, at the elevator exit, to know what is the correct side to take looking at the room number indicators. It is always preferable to look for cues and possible hints using an increased contextual attention before moving. During the elevator ride, your attentional reach cannot be expanded and you dedicate brain power to other matters while being confident that your next subgoal (exiting at the correct floor) will be fulfilled. After having performed the path from the entrance to your room several times, habituation already started to kick in. Sampling minimum information from the environment, you can perform the trajectory nearly blind folded, for instance, only rarely looking at the color of the floor carpet while checking your phone messages. From a Reinforcement Learning (RL) perspective we can model this navigation task with a grid world including all these elements as in Figure **1** (entrance to the hotel in white, lobby in blue, elevator floor in red, corridor in green, your goal/room dark green). The agent (depicted as an isosceles red triangle facing its front direction) can have egocentric actions (go forward, turn left and right). An observation corresponds to a local view around the agent position (depicted as a 3×3 red shaded area around the agent in Figure **1a**. The action space is augmented with an additional action to increase the observation size in a concrete environment that we will use (see Figure **2h**). We make the following analogies in Figure **1a**: the goal cell (marked with a flag) is your hotel room (a terminal state, when reached the trial/episode terminates), the starting positions of the agent (one and two) are the hotel entrances. Then, the lobby (in blue), the elevator (in red), and the room floor (in green) follow. Grey cells are non traversable walls. The two elevator’ rows correspond to the two corridor entrances. The fact that you do not consciously select a row of elevators is modelled as having a random starting position (indicated by 1 and 2 in Figure **1a**). There could be an alternative modelling: adding a blocking intermittent cell at the entrance of the elevator cells. The advantage of using two initial positions instead is that the environment remains stationary and simpler, with better convergence guarantees of Model-Free approaches. As mentioned earlier, previous work consider non-stationary tasks [2, 6, 19, 27] forcing to what we call a ‘‘smoother” arbitration of Goal-Directed and Habituation behavior. As in Two-Choice tasks [6], we can add probabilities of position restarts (establishing a common and rare, in this case, initial transition). We could say that the habituation process manages to make us take the common elevator side with 0.8 probability and that, there is a free and available elevator on the other side with 0.2 probability. Interestingly (and as we will see in the results Section 3.2), the uncertainty in the learning can be controlled by changing the restart probability of the common starting position from 1.0 to 0.5. We can finally set the simplest reward contingency possible in which the agent receives a negative reward at every step (as we want to get to our room as soon as possible).

**Figure 2:**
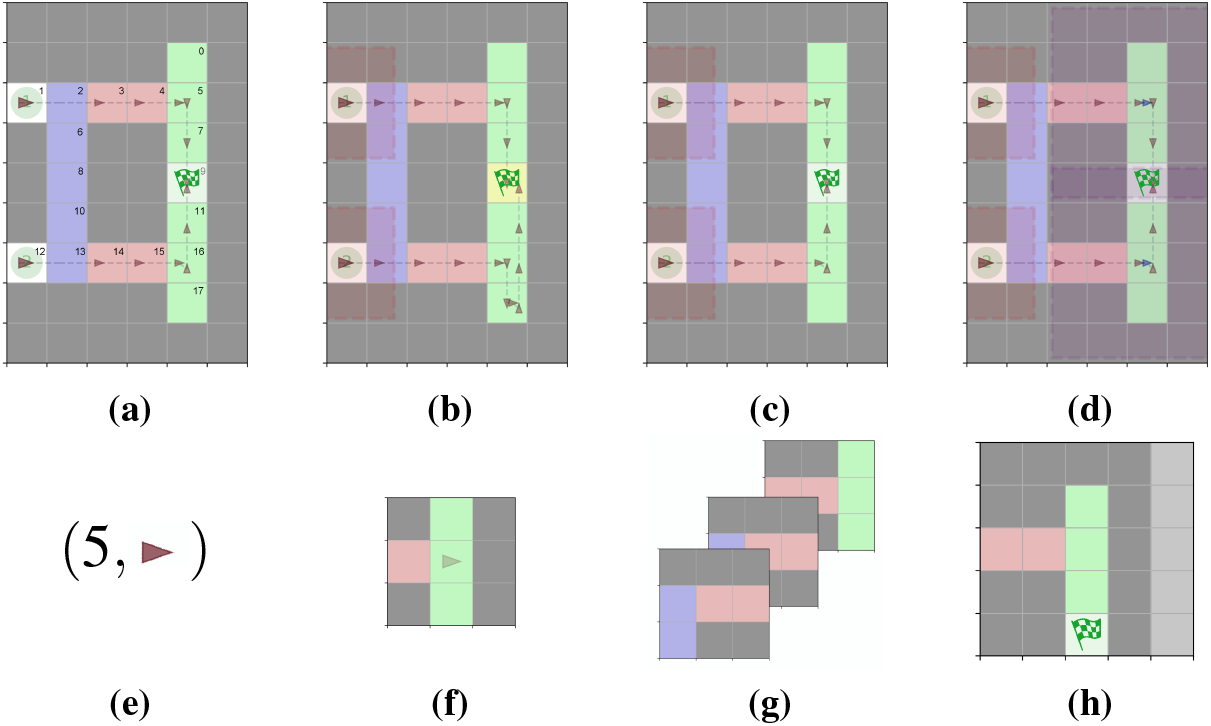
Environments (a,b,c,d) and their associated observations a the so-called bifurcation point (e,f,g,h). **(a)** Global environment grid and its associated observation at the bifurcation point is shown in **(e). (b)** Partial Observable environment and its associated observation at the bifurcation point **(f). (c)** The memory environment and its associated observation at the bifurcation point **(g). (d)** Observation Size On-demand environment and its associated increased observation at the bifurcation point **(h)**.

Using a Model-Free RL approach like Q-Learning, we can learn a policy that maps states to actions to be performed at each situation. Your policy can be something very simple: go forward while seeing the blue carpet of the lobby, enter the red elevator at floor 0 and exit at floor 1. Then, turn right when exiting the elevator at the green carpet and continue forward. But partial observability can pose problems to Model-Free learning. When you take the rare transition (uncommon elevator row), your habitual control brings you to the wrong corridor. But what enables you to detect that you are in the wrong corridor? The brain cannot afford to delegate all the responsibility to an habituation mechanism that can fail. It needs to monitor A possible hypothesis is that while executing an Habitual/Automatic policy, we have a mismatch checker in the background. This mismatch detector can only be Model-Based. You have a kind of an idea of what states you traverse in your habitual path to your room and you have also a sort of timing expectations of when to get there. A Model-Based mismatch detector can be vigilant to possible Habituation failures. This “mismatch detector” fits perfectly with the so called Successor Representations in RL (SRs), which are a kind of relaxed versions of learning the exact transition model of the environment, in which order of state visits is lost. Learning SRs you can have access to which states are expected to be visited after a certain current state.

If your habitual behavior is to take the left row of elevators (top corridor in Figure **1**), then the learned SR will cause a high temporal difference error (being detected as a very unexpected event) when visiting the dead-end after the elevator’s exit. The fact that you find yourself in the wrong corridor is vivid and unexpected. How do intuitively solve this anomaly? and what do we do to prevent this to happen in the future? One possibility is that, next time, we force ourselves to remember from which elevator side we came from. This solution would indeed solve the anomaly, but its costly in terms of allocated resources. But wait, its only one bit of information what is required to remember.. yes, but you need to remember it during all the elevator trip and this may interfere with other thinking processes. And its effectively the case, with sufficient memory of the experienced states, any partial observable problem can be resolved. But this is not how we solve the anomaly.

The hypothesis stated here, is that we have an additional Model-Free mechanism that is an alarm activation that forces a switch from Habitual to Goal-Directed behavior. So, when the SR mismatch detector is surprised with a high TD-error, we do a retrospective Goal-Directed assignment of a warning signal in the precedent state when we made a decision, when we exited the elevator. That state is assigned a warning signal, in such a way that, when we exit the elevator again, we go: but wait… something bad happened here at this particular state. This alarm warning is activated via a Model-Free mechanism that then switches to a Goal-Directed enhanced attentional process in which we widen our context observation, acquiring more state “resolution” in the search for clues to make the correct decision; the right context is added.

### 1.3 Related Work

We describe in the following the state of the art of different computational modeling approaches to Habituation (Model-Free) and Goal-Directed (Model Based) arbitration.

#### Speed/Accuracy trade off

In [18] a measure for state-action value uncertainty is introduced called Value of Perfect Information (VPI). VPI is computed according to the intersection of the Gaussian distributions approximation of the value of the two best actions at a certain state. If VPI is high (if the intersection is high) it is worth losing some time and resolve the uncertainty by planning operations because the second best action may become better than the current maximum value one. The VPI measure is used for arbitration between Model-Free and Model-Based methods. The learned value of decisions (the state,action value of the so-called *Q*-Function) in ambiguous (partially observable) situations (states) oscillates without convergence. This makes the so called Value of Perfect Information (VPI) to be high and increase, no matter how many episodes we experience, or model based queries we perform.

#### Retrospective MB Inference guides MF updates

In [27] a mechanism is modeled and identified behaviorally by Maximum Likelihood model explanation of the behavioral data. In the presence of incomplete information, Model-Based inference of the hidden state or its related states are reinforced positively or negatively.

#### Parallel MF-MB processes

In [30] a similar procedure of fitting of models to behavioral data is used [27], and a parallel execution of Model-Free and Model-Based processes is found to have more predictive accuracy.

#### TD-error based replay and consolidation

(a) Dyna-Q [31] is the prototypical Model-Based algorithm. It interleaves learning from simulated experiences (extracted from the learned model) with usual Model-Free learning. The arbitration is driven by the *TD*-error exceeding some predefined threshold. The intuition would be that when we find a high surprising reward at a certain state, it is worth adding replayed reinforcement updates to th states that lead to the current state. (b) SR-Dyna [28] is based on Dyna-Q and uses Model-Based updates to learn the Successor Representation (SR). SR learning sits in between Model-Free and Model-Based approaches. SRs gather information about future expected revisitation of states given an executed behavioral policy (see for a very recent review [24]) and can be learned through MF TD-Learning. When combined with the actual reward (not discounted) that can be obtained in each state, a value function can be obtained. SR based methods, learn the expected revisitation and state reward separately, thus being more robust and adaptable to changes in the reward contingency [28].

#### Distributed Adaptive Control (DAC) architecture [10]

The Adaptive layer (learning state to action associations) can be considered a Model-Free learning module and the Contextual layer (stores and makes queries among state-action sequences or graphs) can be equated to a Model-Based module. The prototypical DAC task described in [10] is a partial observable problem similar to the one described here. The mechanisms of arbitration in DAC between the Adaptive and Contextual layers are basically two: (a) the contributions of the two layers are summed up in motor space, (b) the Contextual layer overrides Adaptive layer commands.

#### Local/Global context networks

In [8] different state resolutions are used to perform Model Free learning. Local and global versions of the current state are used to inform the MF decision. Local and global derived decisions are added up or selected with an ad hoc mechanism. This previous work opens the path of considering Model-Based decisions as a mechanism of enhanced attentional actions.

#### Meta-Learning

Meta-Learning has flourished recently in various flavors. See for example the review [3], in which the title “Reinforcement Learning: fast and slow” refers to Meta-Learning (fast) and Model-Free approaches (slow) respectively. The idea here is that meta-learning can learn the structure of the task and is faster to adapt to a new variation. This “fast and slow” terminology can be misleading when compared to Kahneman “Thinking Fast and Slow” [17], which as said in the introduction, refers to Habitual and Goal-Directed systems respectively. Meta-Learning can be seen as Hierarchical RL problem where a nested outer loop learns parameters of the inner loop, goals for example [20].

## 2 Material and Methods

The HER anomaly can be modeled as a grid world environment that we will ultimately define as a Partially Observable Markov Decision Process (POMDP). This will allow us to discuss Reinforcement Learning (RL) applications to the task. RL suits perfectly to model habitual behavior: actions that we learn to execute automatically

### 2.1 The environment as a Partially Observable Markov Decision Process (POMDP)

MDPs are useful for modeling control processes (here they will play the role of the environment) in which discrete decisions can be taken (actions). The MDP will describe the possible states and the effect of actions taken on those states. Formally an MDP is a tuple 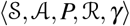 where 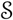 is the set of states, 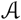 the set of actions, *P* the transition probability distribution 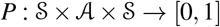 satisfying ∑*_s′_ P*(*s′*|*s, a*) = 1 which defines the Markovian dynamics of the states conditioned on actions of the agent, *R* the expected reward function for every state and action 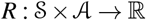 and *γ* the discount factor of the return (reward obtained from state *s* and discounted geometrically by *gamma*): *G_t_ = R*_*t*+1_ + *γR*_*t*+2_ + *γ*^*t*+2^*R*_*t*+3_ +…. The aim of Reinforcement Learning (RL) is to learn an optimal policy that maps states to actions 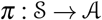 such that future expected reward is maximized: 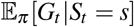 being *S_t_* the state at time *t*.

A Partially Observable MDP (POMDP) is a generalization of an MDP in which some states 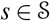 may not be directly observable by the agent. Instead of observing fully identifiable global states, the agent receives the so-called observations 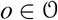 that depend on the current hidden state via an observation process *H*(*o|s*). Then, formally a POMDP is augmented with the observation set 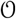 and *H*, being a tuple 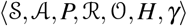. We use the discrete state space version of POMDPs, although continuous versions can be defined as in [33]. In the following we use *s* and *S* for denoting all types of states (observable or not). In fact we will add a superscript to the state notation *S*^*r*=1^ that is indicative of the degree of observability of the agent, as we explain in the following Section 2.2.

### 2.2 Grid-world Environments

We describe the different grid-worlds scenarios that together with the RL algorithms will match the different solutions to the HER task shown in Figure **2**. We tried to minimize the exemplar environment (see Figure **1**) that will accompany us throughout the discussion to a minimal size containing all the interesting elements, and we will further simplify it to a One-Choice task similar to [13].

In terms of the MDP (as described in Section 2.1) common to all variations: we fix the reward contingency as being: −0.5 for every step (independently of the action) and a supplementary +1 for getting to the room. Thus the trajectory depicted in Figure **1b**, starting at entrance 1, has a total reward of −2.5.

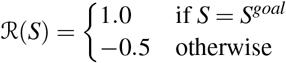

The action set available at every state is 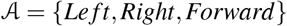 and includes an extra action for the *Observation Size On-demand* environment 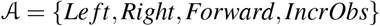, that we call *A^IncrObs^*.

1. **Global**. The first MDP version corresponds to the environment in which the agent has a global view, each state can be linked to an integer identifier and the agent has access both this number and its facing direction that could be identified as well with an integer (from 1 to 4 according to each direction: north, east, south, west). An example of observation at the bifurcation cell (numbered 5) can be seen in Figure **2e**. As numbered in Figure **2a** there is a total of 17 ∗ 4 = 68 states: the number of free cells multiplied by the 4 possible agent directions. The agent has perfect information in this environment at all times.
2. **Partial Observable**. The Partial Observable (PO for short) version of the environment the agent has incomplete information and has only access to a local view around his position, rotated, facing its front position. This default local view that we will use corresponds to a 3×3 matrix centered in the current position of the agent. This observation will be considered of having radius 1 and will be called *S*^*r*=1^ and it does not need to contain agent information as the 3×3 matrix is rotated according to the agent facing direction. In Figure **2b** the observation area is represented by the red shaded area and example visualization can be seen in Figure **2f**. A radius 0 observation, *S*^*r*=0^, corresponds to the state of the current cell, an alternatively one can add the facing orientation of the agent (an integer between 1 and 4), we will call it *S*^*r*=0,+*o*^ adding a +*o* to distinguish it. The observation could as well be linked to an integer identifier and some differentiated global states collapse into the same local partial observable ones. In our example, the agent can observe 60 different partial observations from a total of 68. For instance, at the bifurcations points (as depicted in Figure **1** (were we show all the observations of two trajectories), the agent will have the same observation, and thus will be unable to distinguish its real global position.
3. **Memory**. In this environment we add memory of the last visited states. The observation consists of the current local view and a history of the *n* precedent ones, it is a concatenated version of the last *n* local views of the *n* visited states. We have two options, either the environment deals with the delivery of the observation history, or the agent gathers the observations using a memory buffer; both are equivalent. For example, if the history contains the last 2 observations and the current one, the state would be *S* = 〈*s*_*t*−2_, *s*_*t*−1_, *s_t_*), and the agent would update its Q-Function accordingly. Note that the number of states increases exponentially with the history length. In Figure **2f** we show such an observation at the bifurcation point.
4. **Observation Size On-demand**. The Observation Size On-demand environment subsumes the Partial Observable one and includes an additional action at every state. By default the agent receives observations of radius 1, *S*^*r*=1^, size 3×3. This new action delivers an increased observation size as depicted in Figure **2h** without moving and thus losing a time step, getting the usual negative reward step −0.5. We will call this action: Increased Attentional action. The so-called increased size observation has radius 2, *S*^*r*=2^. The environment becomes completely observable with radius 2 as all states are unique.

### 2.3 *Q*-Learning Model-Free algorithm

*Q*-Learning is the prototypical Model-Free algorithm and its associated with an Habituation mechanism. For learning a policy, Q-Learning doesn’t make use of any model of the environment, thus the denomination Model-Free. It can be compared to an habituation mechanism as at each step we update the state,action according to a reinforcement mechanism that depends on the reward obtained and making a decision is quick and automatic as it requires only access to the pre-computed table: the *Q*-Function. *Q*-Learning solves the RL problem by estimating this state-action value function 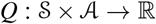 which stores the future discounted reward that can be obtained when being at state taking a certain action. A policy can be obtained by “greedyfication” of the Q-Function, *π*^*^(S) = argmax*_a_Q*(*S, a*), by selecting the highest future discounted value action at each state.

The *Q*-function *Q*(*S, A*) can be stored in a table, the so called tabular form. When the state space is high dimensional as in the case of Atari games where a state 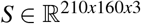 a Deep Convolutional Neural Network (DCNN) is used to approximate *Q* (as in DQN [22]). In this paper we can remain in tabular form without loosing scope, on the contrary, convergence properties hold with the tabular case and not when approximation is used (which introduces uncontrolled partial-observability).

For selecting actions *Q*-Learning uses the *ε*-greedy policy *π^ε^*(*S*), which addresses the exploration-exploitation tradeoff by selecting random actions with probability ε and exploiting the current estimated *Q*-function argmax*_a_Q*(*S, a*) with probability 1 − *ε. ε* starts at 1 in the beginning of learning and goes down to 0 (usually linearly or exponentially).

At every step, when observing state *S*′ and reward *R* after performing action *A* at state *S* (the tuple *S, A, R, S*′) Q-Learning performs the update:

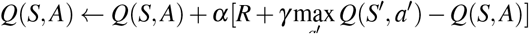

where *α* is the learning rate and *γ* the discount factor. The part *δ = R + γ*max_a′_ *Q*(*S*′, *a*′) − *Q*(*S, A*)) of the *Q*-Update is also called the Temporal Difference error 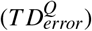 and it has often been related to dopaminergic signals in neuroscience as it can be interpreted as a reward expectancy signal and provides expectations of reward outcomes. For instance, after having observed the environment transition from *S_t_* to *S*_*t*+1_ when executing action *A_t_* and obtaining reward *R_t_*, the 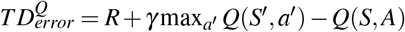 will be close to 0 when that reward was expected (in complete information environments, this will happen typically after learning), negative when it was overestimated (lower than expected) and positive when it was underestimated (higher than expected).

### 2.4 Policy gradient methods

Policy Gradient (PG) methods solve the RL problem directly by computing a policy: not like Q-Learning that estimates the state,action value function and then obtains a policy by the mentioned “greedyfication” step. PG methods also update the policy in the direction of the gradient. This is also a distinction with value based methods which updates the Q-function state,action value proportionally to the error positively or negatively reinforcing the action taken in the corresponding state (rule with several names: decorrelation learning, delta rule…). The latter being simpler, has been more related to neuroscience principles. The simplest and prototypical PG algorithm reinforce is a Montecarlo based method, meaning that it updates the policy from a complete episode (also called rollout).

PG methods can deal with stochastic policies which would allow for different solutions to partial observability. For example one could think of acting random when exiting the elevator. As we mention in the results Section 3.2, the Reinforce algorithm is not able to learn an optimal stochastic policy in the case of the HER anomaly.

### 2.5 Dyna-*Q* Model-Based algorithm

The Dyna-*Q* is the prototypical Model-Based algorithm and it builds on *Q*-Learning, adding *Q*-updates that do not come from real experience (*S, A, R, S*′ tuples obtained from the environment), but come from a learned model; that we learn simultaneously as we estimate the Q-Function. That is, at every step, when observing the tuple *S, A, R, S*′ we can learn the model transition *Model*(*S, A*) ← *S*′, *R* and its reversed version *Model_bck_*(*S*′) ← *S, A, R*. If the environment (the MDP) is deterministic we can just store all transitions in a directed graph. If the environment is non-deterministic we need to store and maintain the probabilities of every transition.

If the TD error, 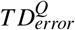, at the current step is greater than a threshold, we make random *Q*-updates from previously observed states and actions taken from the learned model. This model updates are considered as a planing contribution to the estimation of the *Q*-Function and make use of a learned model, thus the term Model-Based.

Dyna-Q model updates are reminiscent to the experience replay updates in DQN algorithm, used to make the training examples independent and identically distributed (i.i.d.) [22]. Updates can be sorted by priority (for example using increased 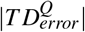) as described in [32] and used in [21]. Sorting updates by *T*D-error has also been used in experience replay techniques.

Putting together sorted 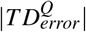 and backward state model updates, leads to the enhanced version of Dyna-Q called Prioritized Sweeping, that includes model updates of the state predecessors to the current one when the 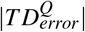 is greater than a considered threshold. This version is the one used in [21] to explain several neuro-physiological results related to forward and backward hippocampal cell firing.

Its important to notice with Model-Based Dyna-*Q* approaches that there are updates to the value function *Q*(*S, A*) that come from real experience and others come from simulated one extracted from the recorded ones in the model.

### 2.6 Successor Representation learning: in between Model-Free and Model-Based

Successor Representation (SR) consist in learning a relaxed version of a state transition model (as in Dyna-*Q*). Instead of learning and maintain a full transition directed graph (or the MDP itself) we learn the future discounted probability of revisitation of every state from every other state. The order in which future states will be visited is lost. For this reason we are going to refer to SR as a Semi Model Based method. This discounted expectancy is computed in the matrix *M*(*S, S*′) that maintains the discounted visitation probability that *S’* will be visited after being in S. The *M* matrix can be learned as in *Q*-Learning by Temporal Difference updates. The matrix *M* must be initialized with the identity: *M*(*S, S*) = 1. *M* can be learned online by storing its adjacency list. When a new state is encountered it must be initialized by adding a new row with a 1 in its column id and a 0 in all other rows. After observing the environment transition from *S_t_* to *S*_*t*+1_ the SR estimate *M*(*S_t_, S*′) is updated for all encountered states *S*′:

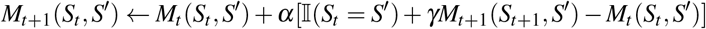

where *α* is the learning rate, 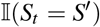 is equal to 1 if *S_t_* is equal to *S*′ and 0 otherwise. The Temporal Difference error, in this case 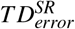, is the part of the update that multiplies the learning rate 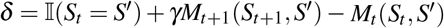. The 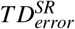 indicates how surprising is to encounter *S*′ after *S_t_*. After experiencing the environment transition *S_t_ → S*_*t*+1_, we have *S*′ = *S*_*t*+1_ and the 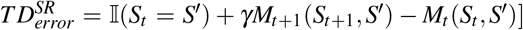 simplifies to *γ-M_t_*(*S_t_*,*S*_*t*+1_). If the TD_*error*_ is positive it means we are underestimating *S*_*t*+1_ visitation after *S_t_* and *M*(*S_t_*, *S*_*t*+1_) should be increased; if it is negative, it means we were overestimating it, and *M*(*S_t_, S*_*t*+1_) should be decreased. So we can take the absolute value for a surprise signal: 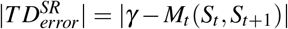. If |*TD_error_*| is close to 0 would mean that we are not surprised of having encountered *S*_*t*+1_ after having visited *S_t_*.

We are then going to use the SR updates inside *Q*-Learning algorithm. While we learn to habituate with Q-Learning, SR updates will compute the future discounted expected visitations. That is, after experiencing a real environment transition *S_t_ → S*_*t*+1_ during *Q*-Learning, *M*(*S_t_, S*′) is updated for all states *S*′.

After a policy *π*(*S*) is learned with Q-Learning, and we are exploiting it at state *S*_*t*−1_, executing action *π*(*S_t_*), the environment transitions to *S_t_*. At each environment transition a surprise signal can be obtained by remembering the previous visited state 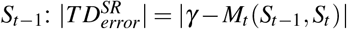. We are providing results of this surprise signal under partial observability in Section 3.8.

Lets discuss about the effects of *γ* in the learning as done in [24] and [29]. When *γ* = 0 the initial identity matrix is obtained. When *γ* is close to 0 we obtain the identity matrix plus some faint presence of the real one step transition matrix in which order information is present. When *γ* is close to 1 we obtain the infinite discounted visitation expectancy matrix in which order is lost. After learning, usually 0 ≤ *M*(*S_t_,, S*′) ≤ 1, but it can happen that *M*(*S_t_,, S*′) > 1 when there are states that can be visited several times in the same episode. This is not normal when there is complete information, but it can happen often in the presence of partial observability like in the HER task.

We are not using it in this paper, but the value function can then be computed from the matrix *M* and the mean reward obtained in a state. To approximate the state, action value function *Q*(*S, A*) one needs to extend the definition of the SR matrix to *M*(*S_t_, A_t_, S′, A′*) being the future discounted expectancy to observe and perform state, action *S′, A′* after having been in state *S_t_* and having performed action *A_t_*. Learning these two quantities separately give advantages when the environment is changing (non-stationary) as relearning is quicker. These gives rise to variations of SR driven learning [28] like SR-Dyna, in which SRs are learned via model updates as well, as in DynaQ. The HER task is stationary and an SR learning approach would not perform better.

Successor Representations have been found to be used in the brain, which would be expected being relaxed versions of exact transition models. Maintaining a structure that relaxes order seems also beneficial as its less costly. Sometimes you do not know about the exact order in which you should expect states/features but all of a sudden you are aware of an state that it was not expected. We will use this fact when discussing about being aware of the HER anomaly when it actually happens, and you find your self in the wrong corridor. We hypothesize that an SR is being used.

SRs can also provide expectations of timely events (state visitations) [24, 25]. Imagine in the HER anomaly that the wrong corridor for finding your room was longer. A brain signal could be warning you that your goal was usually found before in time. SRs could be as well used for that purpose.

## 3 Results

### 3.1 Reduction to a Two and One-choice task

The HER task and its RL formulation can be reduced to a One-Choice task (see Figure **1c** as in the partial observable One-Choice task in [27]. In fact the first Two-Choice task that originated a tree of research papers, is in fact said to be more adequate to be translated to a One-Choice in [2].

We take some time to explain in detail the task in [27] as is the only one considering incomplete partial information. In their case, when the participant faces the so-called *uncertainty trial*, you are randomly re-spawn to one of two states, combinations of object pairs (*s*_1_ = {key/telephone, stove/bulb}, *s*_2_ = {key/stove, telephone/bulb}), and you make a one choice selection of which pair of objects to chose from the state in which you started. Then the environment (the oracle in the paper) makes a random selection (hidden to you) of an object from the pair you chose and you are given feedback of the reward you got (related to the object the oracle chose) and the room associated to that object (which is in fact associated with another object as well). The reward associated to each object is non-stationary, changing in time. In [27] a retrospective inference is proven to be present that infers which object was chosen by the oracle according to the feedback of the room it was provided and reinforces (positively or negatively) according to the reward that was received.

In our case (see Figure **1c**), there is as well one single decision to make, to go left or right in the middle state (exit of the elevator), and the success of that action will depend on from we came from (the state we started from). In the case of the *Observation Size On-demand* environment, there is an extra action in the middle state that makes the environment completely observable.

The reduction to a One-Choice task improves the sample efficiency of the algorithms, but does not affect any of the results that we present in the following sections: the anomaly is present, it can be solved with memory methods, DynaQ fails on this task due to partial observability, and retrospective inference can be performed to raise an alarm signal at the decision point to enhance attention and make the task completely observable.

Moreover we preferred to realize all the experiments with the designed environment although many of the available actions (3 or 4 actions at every state) do not correspond to the real available actions when solving the navigation task to your hotel room. Many actions are discarded from the beginning by our previous knowledge of having interacted with walls, corridors and elevators. But doing so we could fine tune much more adequately the action value uncertainty measures that we define and test in next Section 3.3.

### 3.2 Habituation / Model Free evaluation on the HER task

We present Q-Learning results, the prototypical Model Free habituation mechanism. We first focus on its evaluation on the Global and Partially Observable environments. The Global environment (see Section 2.2) avoids the HER anomaly as every state is unique and a correct turning action can be learned at each crucial decision point (when exiting the elevator), to the eyes of the agent, each decision point is a different state. These global states are the ones used in all studies [6, 28] except the recent [33] (which considers noisy uncertain observations, another aspect of partial observability) and the previously mentioned [27].

The HER anomaly is reproduced in the Partial Observable environment. What do we mean by reproducing the HER anomaly? We mean that, after learning (and no matter how long training lasts), one of the trajectories starting at each initial position always includes an undesired detour due to an habituation failure (see Figure **1** for an illustration of the two trajectories and Figure **2b** in which it is put in context with the other environments optimal trajectories). The HER anomaly is reproduced only when considering the right level of partial observability. In our case this corresponds to a state size 3 × 3, or as we described to a state observation having radius 1, *S*^*r*=1^. In this situation, the states at the two decision points collapse into one (in fact, a total of 68 states collapse into 60). A radius of 2, *S*^*r*=2^, makes the task completely observable again. Reducing the radius to 0 (and the agent only perceives the cell it is in) makes the task unlearnable. When having the right level of partial observability (states *S*^*r*=1^ having radius 1), the anomaly can be reproduced in fact by all MF (Q-Learning), MB (DynaQ) and in-between approaches (SR-Dyna [28]). Moreover, in the MB standard DynaQ, the effect of partial observability degrades the algorithm performance below Q-Learning. We show the reason for this in Section 3.6.

In Figure **3** we compare the learning performances of algorithms and all environments. Learning performance is assessed (like in [31]) by evaluating the policy 10 times (5 restarting from position 1, and 5 from position 2, see Figure **1**) with no exploration (in the case of Q-Learning based algorithms corresponds to ε = 0) every certain number of episodes of learning.

**Figure 3:**
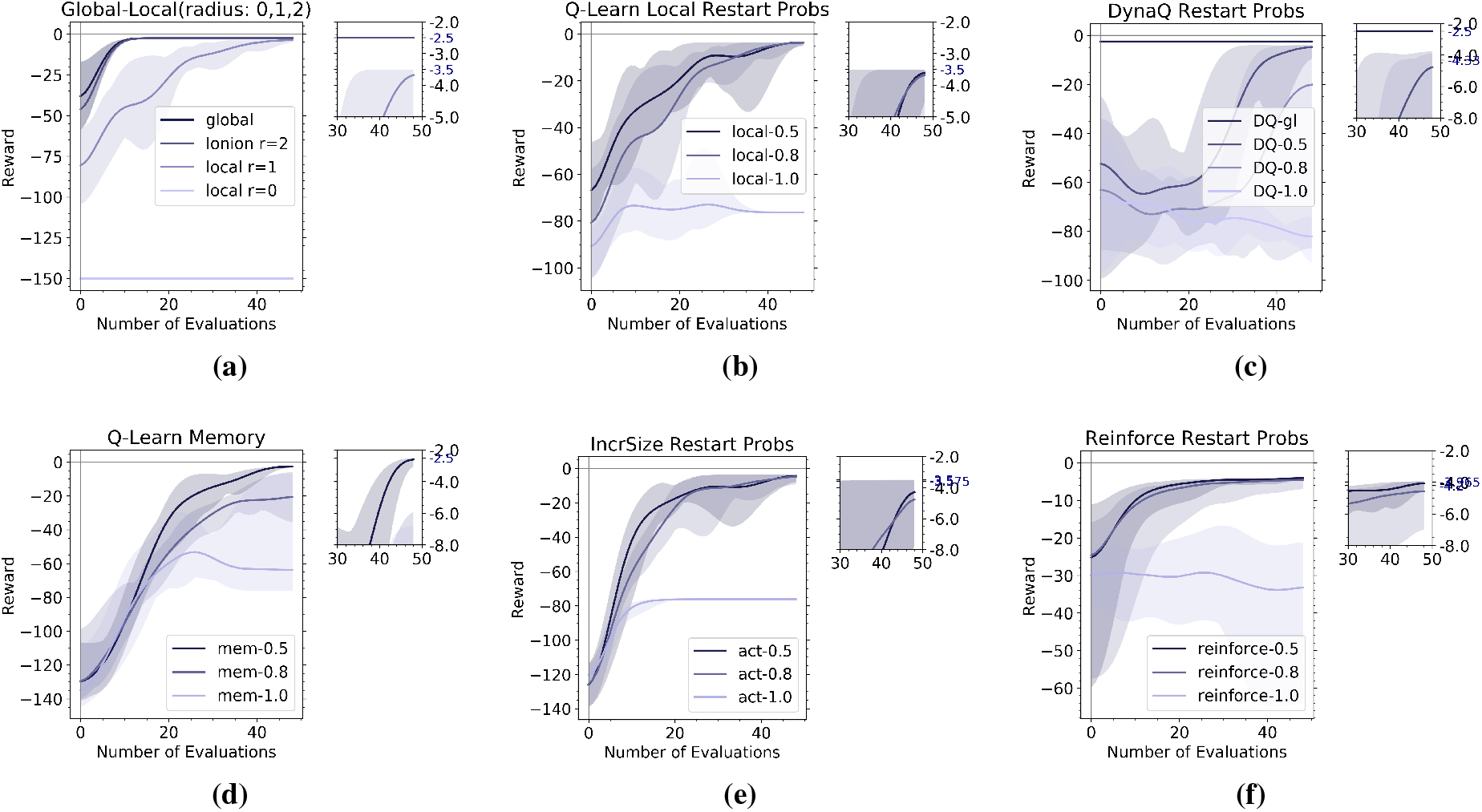
Learning Evaluations.of all environments and all employed algorithms. The mean and standard deviation of the obtained reward for each evaluation is shown for a learning of 1000 episodes (except the memory environment which is executed for 3500), and a zoom of the final evaluation is shown on the right to be able to assess the maximum reward achieved. **(a)** Q-Learning is evaluated on the global and partial observable environments. **(b)** Q-Learning evaluation on the partial observable environment with radius 1, varying the probability of restarting from position 1 of the grid. **(c)** Model-Based prioritized sweeping DynaQ algorithm is evaluated for global and partial observable states, achieving the best performance on the former and a worst performance compared to Q-Learning on the latter. **(d)** The memory environment evaluations with Q-Learning. **(e)** Evaluations of the Observation Size On-Demand environment with Q-Learning. **(f)** Evaluations of policy gradient algorithm Reinforce.

In Figure **3a**, Q-Learning is evaluated on the global and partial observable environments, the latter with observation radius 2 (size 5×5), 1 (size 3×3), and 0 (size 1×1). Radius 2 is equivalent to global states and achieves the maximum possible reward −2.5 (7 steps obtaining −0.5 reward plus the extra termination plus +1). Radius 1 achieves the optimal reward of −2.5 at 5 evaluations, but the other 5 suffer the HER anomaly, a detour of 4 steps, −4.5, giving a mean of −3.5. Having an observation size of radius 0 makes the task unlearnable.

As mentioned earlier, the anomaly could have been modeled with a non-stationary environment by having blinking obstacles (doors) at the entrance of each elevator. We find to be a more simple and elegant solution to use two restart positions and investigate the effect of the restart probabilities, maintaining stationarity. For instance, if starting position one is more probable, then the anomaly is always observed when experiencing the rare transition, that is, starting from position two. We then compare learning performance with varying probability of restarts (1.0, 0.8, and 0.5)at positions for all algorithms and environments.

In Figure **3b** we plot the Q-Learning evaluation on the partial observable environment with radius 1, varying the probability of restarting from position 1 of the grid. For probability of restart 1.0 it happens that the policy evaluation fails in the 5 restarts from position 2 and gets stuck achieving the maximum number of steps which is fixed to 300: thus getting a mean reward of ()5 * (−2.5 + −0.5 * 300) = 76.2). This happens in fact in all of the following cases regarding probability of restart 1.0. For probabilities of restart 0.5 and 0.8, the optimal reward for local observation is obtained: −3.5.

In Figure **3c** show that Model-Based prioritized sweeping DynaQ has the best performance on global states achieving the optimal cost −2.5 on the evaluated policies early on the first evaluations. Nevertheless it is not able to converge properly to find the optimal cost for partial observable environments and has a worst performance than Q-Learning. We comment on the reasons why this happens in Section 3.6.

**Figure 5:**
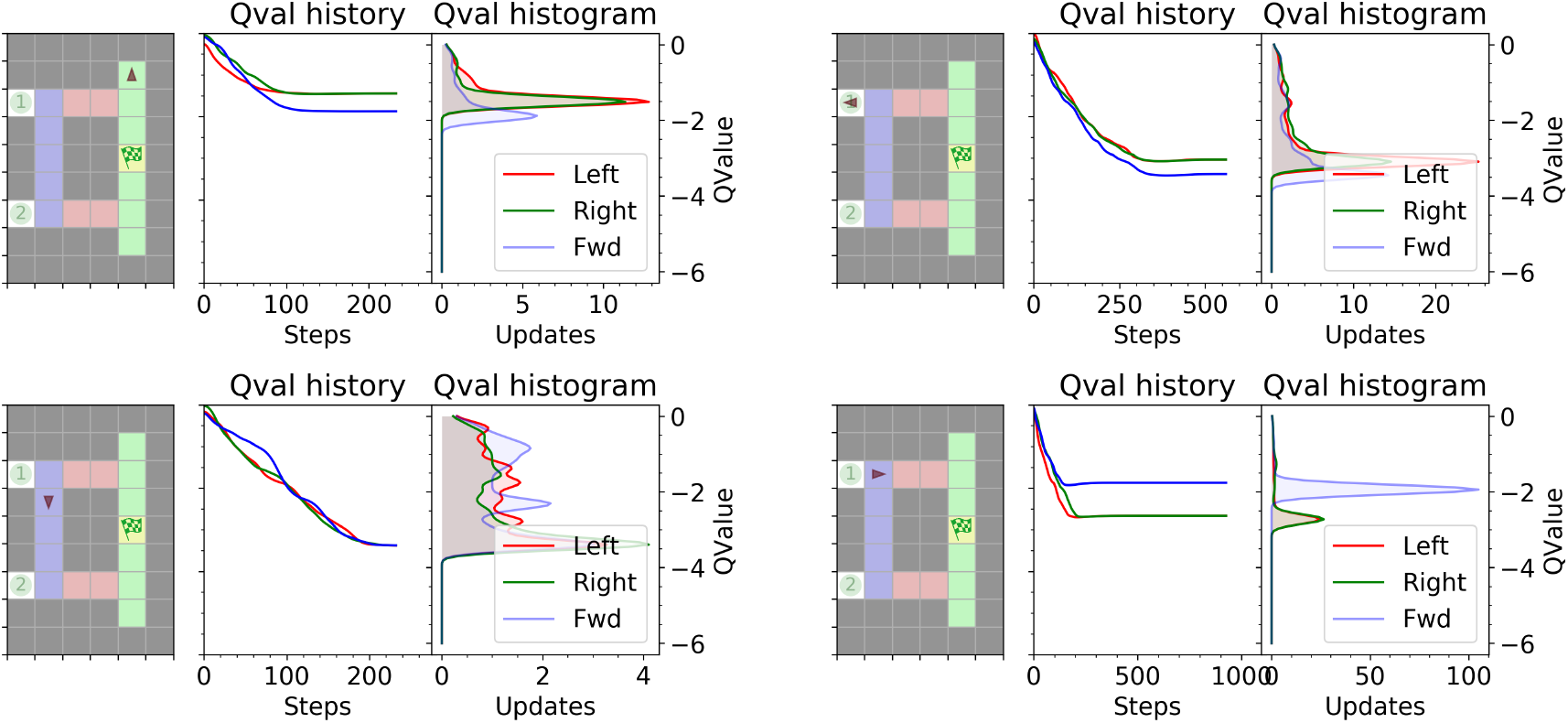
Examples of global states Q-Values update history. We show the history and histograms of Q-Values updates for four different states (explained in Figure **4**). Top left: High *Max2Entropy* as the two best actions were selected equally. In this case the uncertainty comes from the fact that we do not care if selecting right or left as they leave the agent at the same distance to the goal. Top right: Bottom left: The three actions here were selected indistinctly. *Max2Entropy* is still high. Bottom right: Low *Max2Entropy* as the forward action was selected much more. Among the second and third best actions there would be a high entropy.

Figure **3d** evaluates the memory environment and, as it delivers a bigger observation size, needs much more episodes to learn. For this case, we only wanted to show that it can achieve the optimal cost so we run Q-Learning with memory of the last 3 observations for 3500 episodes. The optimal cost of the global environment is achieved for probability of restart 0.5.

Figure **3e** evaluates on the environment with observation size on-demand (an additional RL action is added that loses one step but delivers an increased size observation). One would intuitively think that Q-Learning would find the optimal solution of cost −3, losing one step in the crucial bifurcation point to enlarge you observation size. Only the optimal solution for the partial observable environment is recovered having mean cost −3.5. The explanation for this is dicussed in the following Section 3.5.

For completeness, we also show evaluation results on prototypical policy gradient algorithm Reinforce which does not learn a stochastic policy and converging to a deterministic one having the same result than Q-Learning, but with a worst sample efficiency.

### 3.3 Action value uncertainty measure

We take a deeper look in this section at the action value measures (inline with the Value of Perfect Information (VPI) introduced in [18]). Q-Learning (described in Section 2.3) does at every episode step, an update of the state and action that was just tried, updating the so called value function entry *Q*(*S, A*). Lets take a look at the history of these update profiles, first, in the case of complete observability and perfect information. In Figure **4** we show 5 results plots of Q-Learning with global states / completely observable states.

**Figure 4:**
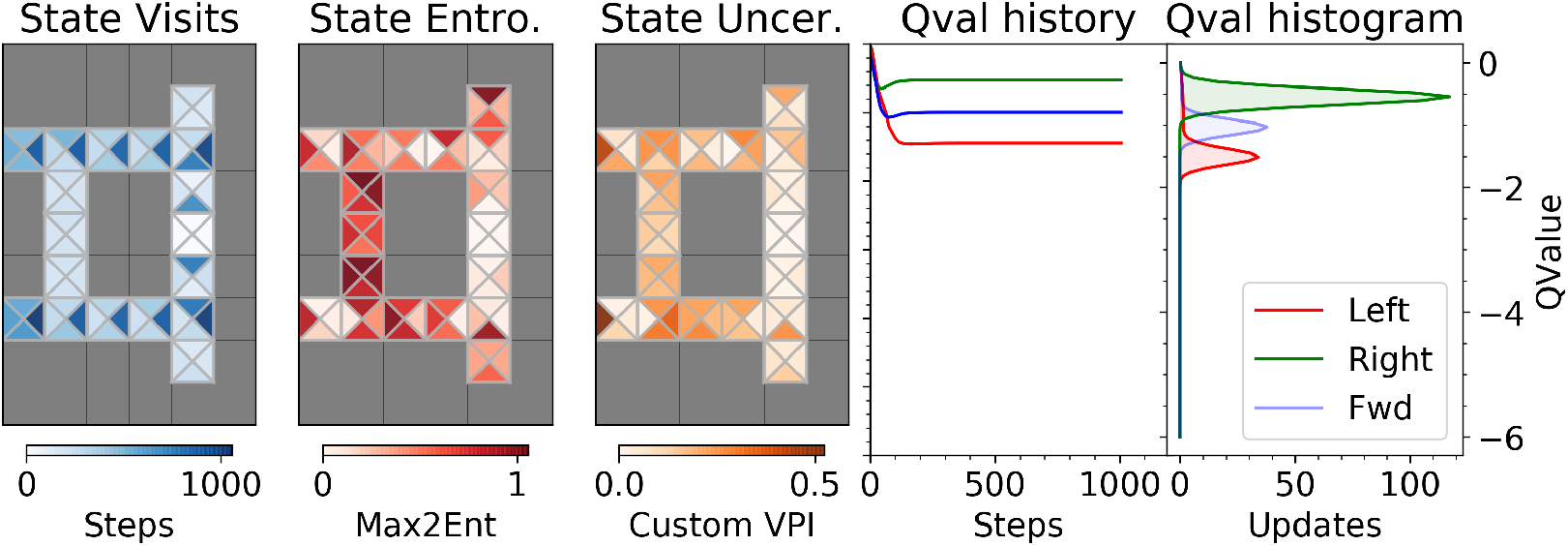
Learning under complete information/observability. First grid plot shows the number of visits that every state received (State Visits). A state here is a combination of the grid position, subdivided with four triangular cells according to the four possible agent facing directions. Second grid plot (State Entropy) shows the entropy measure that we introduce in Section 3.3 for every state. Third grid plot (State Uncertainty) shows the same entropy measure weighted by the visitation ratio of first plot. Fourth plot (Q-Value History) shows the history of Q-values at every step for every action for a particular state (the one that we call bifurcation point, with its associated observations being shown in Figure **2**). Fifth plot (Q-Value Histogram) shows the histogram of Q-value updates. If we do not say the contrary, Q-Value history and histogram plots will always refer to the bifurcation point state.

Lets focus first on the updates that happen on an important state, the so-called bifurcation point (last two columns of Figure **4**). The rewards delivered by the environment at every step are negative (−0.5) except for the final terminal step before reaching the goal, in which the agents gets +1. *Q*(*S, A*) entries are initialized to 0. In the case of complete information the optimal cost is −2.5, thus the optimization is called optimistic because it starts with a higher value than what the optimal can reach, promoting exploration. As seen in Figure **4** (Q-Value history plot) the updates for all actions *a* ∈ {*Left, Right, Forward*} decrease during exploration (high *ε*) and settle afterwards.

We can now compute a measure (similar to VPI) from the Q-Value history plot and histogram, the last two columns of Figure **4**. Lets consider the number of times an action was selected at the bifurcation state. We can see in the histogram of Figure **4** that action “right” was selected around 100 times (because we updated it 100 times) and the second best action was selected around 40 times. Then computing the entropy of the probability distribution of the best two actions, gives us a degree of uncertainty in their selection (what in [18] is computed with the area under the intersection curve of a Gaussian approximation of the distribution). This is exactly the measure that we call *Max2Entropy* and we plot in the grid of the second column of Figure **4** for all states. Uncertainty among the best two actions is highest at several states, not necessarily the ones in which decisions matter. To illustrate this we show the Q-Value history of high *Max2Entropy* states, different from the bifurcation point state in Figure **5**.

If we now take into account partial observability, we observe that the updates profiles of states affected by partial observability have a pronounced variability and are oscillatory (see Figure **6** and Figure **7**). This happens because a partial observable state may have different real states associated to it. This happens even when we set a different probability of restart from position 1 or 2 during learning. We show this in Figure **6**.

**Figure 6:**
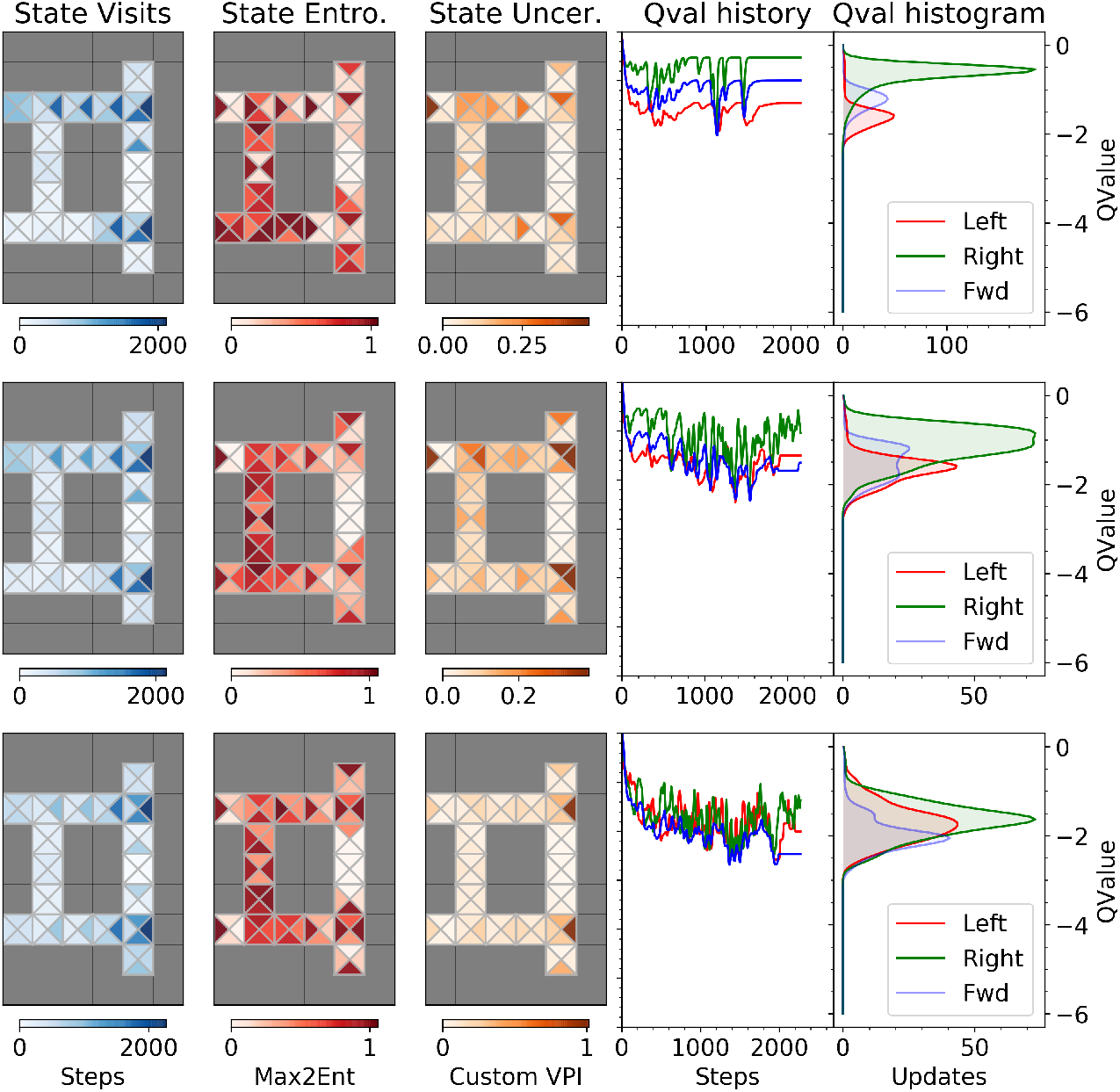
Learning under partial observability. Particularly, we look here at the effect of probability of initial position restart on learning. Columns correspond to the same previously explained in Figure **4**. Now we consider learning under partial observability, the local environment with radius of observation equal to 1 (3×3 size). Each row corresponds to a different learning execution with a different probability of restart from position 1: probability that we restart from the top cell 1.0 (First Row), 0.8 (Second Row) and 0.5 (Third Row).

**Figure 7:**
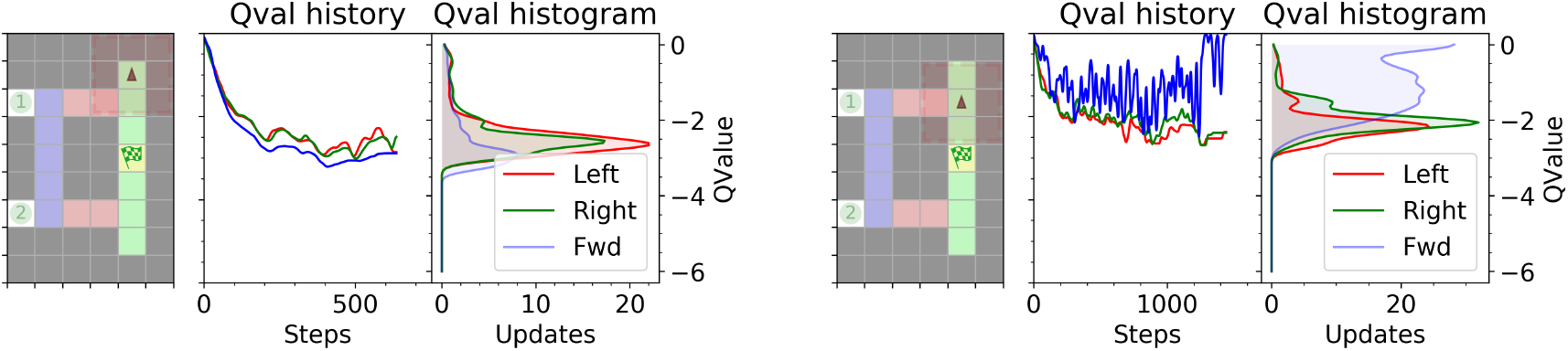
Examples of local states Q-Values update history. Left: Corresponds to the same plot of the global state in Figure **5** (Top Left). The history profile is similar with a high *Max2Entropy* as the two best actions were selected equally. Right: At a partial observable state the value updates can oscillate much more. The most selected action is “forward” and has a low *Max2Entropy*.

### 3.4 Memory approaches to partial observability

The anomaly does not occur when we employ memory of the performed actions or observed states. This result is known: with sufficient memory any partial observable setting can be turned into totally observable.

Figure **2c** with the corresponding observation at the bifurcation point, Figure **2g** (containing memory of the precedent two steps). One can see that, in the designed environment, if the agent has access to at least three history observations, it can disambiguate the situation by knowing from which side it comes from. This is called the “window approach” in [23] which concatenates the most recent *n* observations and treats this as a new state.

Usually memory approaches are used to address partial observability: [23] cites state history recording, recurrency and Neural Turing Machine approaches. Here we propose a new point of view. We hypothesize that, behaviorally, we do not solve the anomaly with a memory strategy. Maintaining in memory from which elevator row/side you came from is costly and must allocate resources during a precious time that one can use for other matters. Thus we propose an alternative way, not described before of dealing with partial observability with an increased attentional mechanism activated “on-the-fly”.

### 3.5 Observation size on demand, adding an RL action

In our first trip to the hotel room we pay more attention and we progressively reduce it as we become habituated to that task. When we experience the anomaly we increase our attentional span again. Attention in this case, is a costly process, to acquire more information we need more time. How can we model this with Reinforcement Learning? To better match this situation we designed in Section 2.2 a new RL environment, the Observation Size On-demand, adding an RL action to deliver an increased observation size, losing one step (reward −0.5). The agent can execute this action at every step/state and receive a higher resolution (increased context) observation (see Figure **2d** and an observation Figure **2h**) at the cost of loosing one step. In previous result Section 3.2 we showed how Q-Learning is not able to learn the optimal policy with this environment: go forward till the bifurcation point is reached, increase the observation size, and then decide left or right accordingly. The Model Free and Model Based algorithms considered here use the same mechanisms for exploration: they start with random actions. Results of Figure **3e** show that the optimal cost policy cannot be learned as it is not easy to learn when to apply the correct increased context action at the right place by applying random actions. We observed that some combinations of running episodes did learn the optimal cost policy, but it is rare. Q-Learning tends to apply redundant attentional actions at useless states, getting a final higher cost, or extinguish all attentional actions getting the higher −3.5 (when −3 could theoretically be achieved). Increasing the learning episodes the latter solution is the most probable.

### 3.6 Goal-Directed / Model Based evaluation on the HER task: Dyna-Q failures

The prototypical Model-Based algorithm, DynaQ is the best performing in the global state environment (denoted “DQ-gl” in Figure **3c**): its evaluations obtain the optimal cost (−2.5) from the very beginning. We can see that this is not the case for Q-Learning in 3a (see legend plot noted ‘global” and “lonion r=2”). In the presence of partial observability DynaQ performs worst than Q-Learning.

The Q-value update history at the bifurcation state (see Figure **6**) oscillate when Q-Learning is still in an exploration phase (*ε* >> 0). DynaQ adds model updates that, in the case of partial observability, increase the oscillation and worsens the convergence even in the case of a common/uncommon (0.8 versus 0.2) restart probability. That’s because model updates are performed independently of any restart probability. In the case of Prioritized Sweeping DynaQ (described in Section 2.5) which adds updates sorted by decreasing 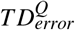 and backward state model updates, convergence degrades even more, as when experiencing rare transitions (like using the non-common elevator), *TD*-errors can reach higher values and useless model updates are added that increase variability and oscillation of the Q-values.

This fact can be clearly seen in Figure **9** comparing the history of Q-value updates for DynaQ in global states (Figure **9a**) and local/partial observable states (Figure **9b**). It is also curios to observe how updates coming from the model are present in the beginning of learning and extinguish towards the end in the case of global states (Figure **9a**) and in the presence of partial observability (Figure **9b**) model updates are present all the time and represent a very high percentage of all updates.

**Figure 8:**
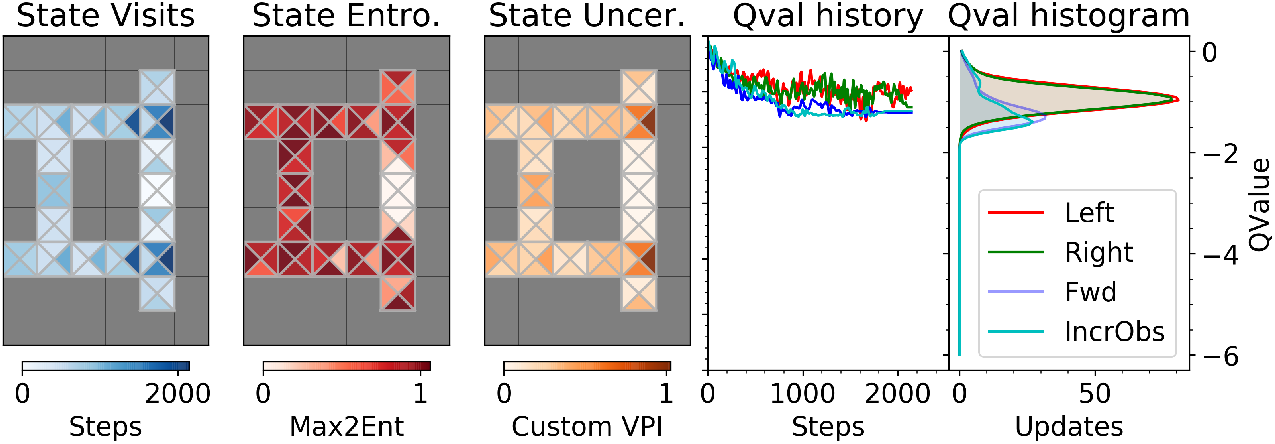
Observation Size On-demand environment with Q-Learning. Columns correspond to the same previously explained in Figure **4**. The increasing observation size action *A^IncrObs^* is tried randomly on every state and creates a much higher *Max2Entropy* map. Q-Learning does not converge usually to the optimal solution of only asking for bigger observation at the crucial bifurcation point. In the histogram we can see that *A^IncrObs^* is selected a few times. Our custom VPI measure is in fact maximum at the bifurcation point.

**Figure 9:**
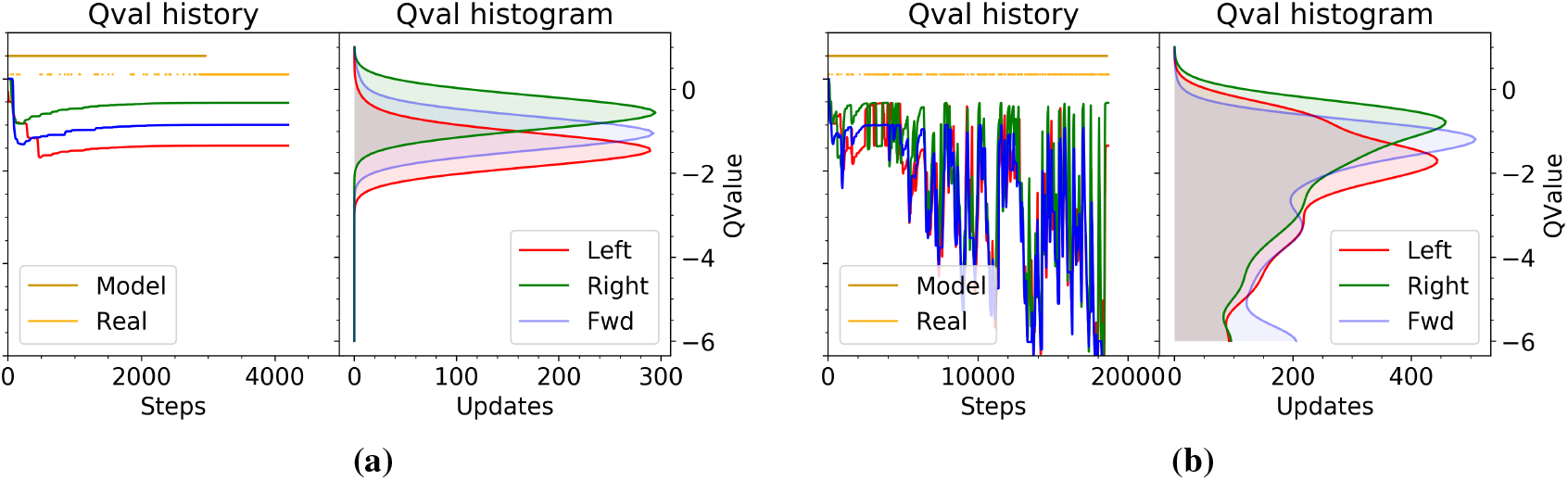
Model-Based DynaQ failures. The Q-Value history and histogram is plotted for the so-called bifurcation state (when exiting the elevator) for Model-Based DynaQ algorithm in the case of global states (Left) and partially observable states (Right). We also superimpose the proportion of Model and Real updates of DynaQ in both cases.

### 3.7 Progressive reduction of the attentional-context: Onion updates

We started describing in the introduction Subsection 1.1 the Goal-Directed sub-goal reaching process that drives the first visit to the room of the hotel: find the elevator, go to the correct floor, look carefully at the elevator’s exit for the correct numbering, etc. We need to find an analogous of this process for the modeling part. We propose that this initial Goal-Directed behavior can be compared to a Model-Free Q-Learning with maximum attention context (radius of the observation size) including learning updates of all sub-observation sizes. The reason for this is that the first time we navigate to our room we allow ourselves maximum attention and we do not care much about time. While paying maximum attention, we also learn about sub details and features of states we traverse. For this reason, we propose what we call Onion updates. An Onion update of state *S*^*r*=2^, means we also update the Q-Function with the states corresponding to the smaller radius *S*^*r*=2^, *S*^*r*=1^, *S*^*r*=0^. We are using tabular implementations, thus it is no problem to perform updates of states of different sizes. If we were using Neural Networks approximations we could pad the reduced states with zeros, and maintain the same observation size (as we did in [8]).

Also Model-Free Q-Learning with onion updates helps acquiring a-priori knowledge about goals. Imagine for example that the goal hotel room is present in the *S*^*r*=1^ observation, correct actions towards the goal can be learned from an *S*^*r*=2^ observation thanks to the onion updates. Also when we progressively reduce our attentional context (reducing observation radius), we do not need to relearn from scratch bigger observations.

In Figure **10** we show the Q-value updates history in the bifurcation point for the Onion Q-Learning execution for the bifurcation state of radius 2 (top row), and bifurcation state of radius 1 (bottom row). In the bottom row one can appreciate the oscillation typical of partial observability that is more prominent than the updates of Q-Learning executed directly with radius 1.

**Figure 10:**
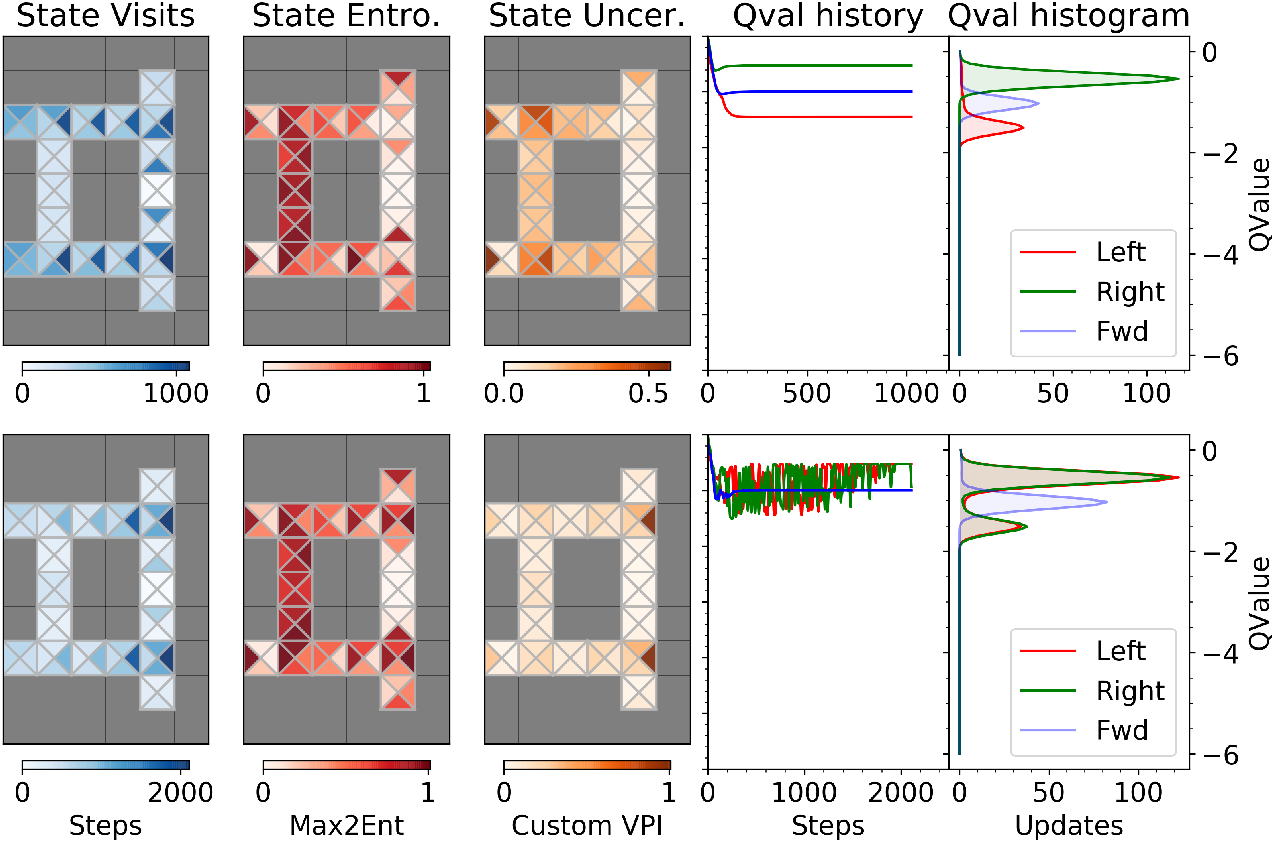
Onion updates learning. Columns correspond to the same previously explained in Figure **4**. Learning with onion updates using states of radius 2, *S*^*r*=2^, makes the problem completely observable. Both rows correspond to the same learning but display different states. Top row corresponds to *S*^*r*=2^. Bottom row corresponds to *S*^*r*=1^.

So here follows a proposal of how we deal with the attentional context. We start with a big context (observation resolution). Add *Q* updates of all inner resolution sizes (onion update). We then progressively reduce the onion-radius of the observations without knowing that in some occasions this will lead to a partial observable trap as in the HER anomaly. If we first start learning from big context sizes, then when a bigger observation will be received (queried) it will serve as an oracle and always will have the correct actions future discounted values associated to it. We provide results of Q-Learning with onion updates in Section 3.2.

### 3.8 Semi Model-Based Habituation monitoring based on Successor Representations

First we are going to discuss about a peculiar moment, the one in which we find ourselves in the wrong corridor. Its a surprising moment in which we realize we are in the wrong place, the eureka moment in which we experience the HER anomaly. This surprising moment could come from the Model-Free mechanism which was driving our habituation behavior; but we argue it is not a reward expectancy signal (the 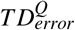 of Q-Learning): we are clearly aware of the fact that we are not in the state we were supposed to be. This surprise signal could as well come from a Model-Based process (or semi-MB) as in Successor Representations, see Section 2.6). Here goes the first hypothesis: while learning to habituate there is a parallel Model-Based mechanism that is checking for mismatches, and possible habituation failures. Successor Representation Learning has been found to be a plausible behavioral mechanism used by humans [26] and could explain a number of rodent electrophysiology studies [29]. We hypothesize that this mechanism is driven by an SR learning that can deliver a signal indicating how surprising is to end up in a certain state after another one. We also hypothesize that a timely SR surprise signal can deliver expectations about the timing in which states are supposed to be encountered. In [25] such a mechanism is described.

In Figure **11** we plot the Successor Representation (SR) matrix *M* computed during Q-Learning (at the beginning and at the end) and a discount factor *γ^SR^* = 0.9. The SR matrix values *M*(*S_t_, S_t_*) can exceed 1 when a state is expected to be encountered more than once in an episode. This situation happens more often under partial observability as several global states collapse into the same local one.

**Figure 11:**
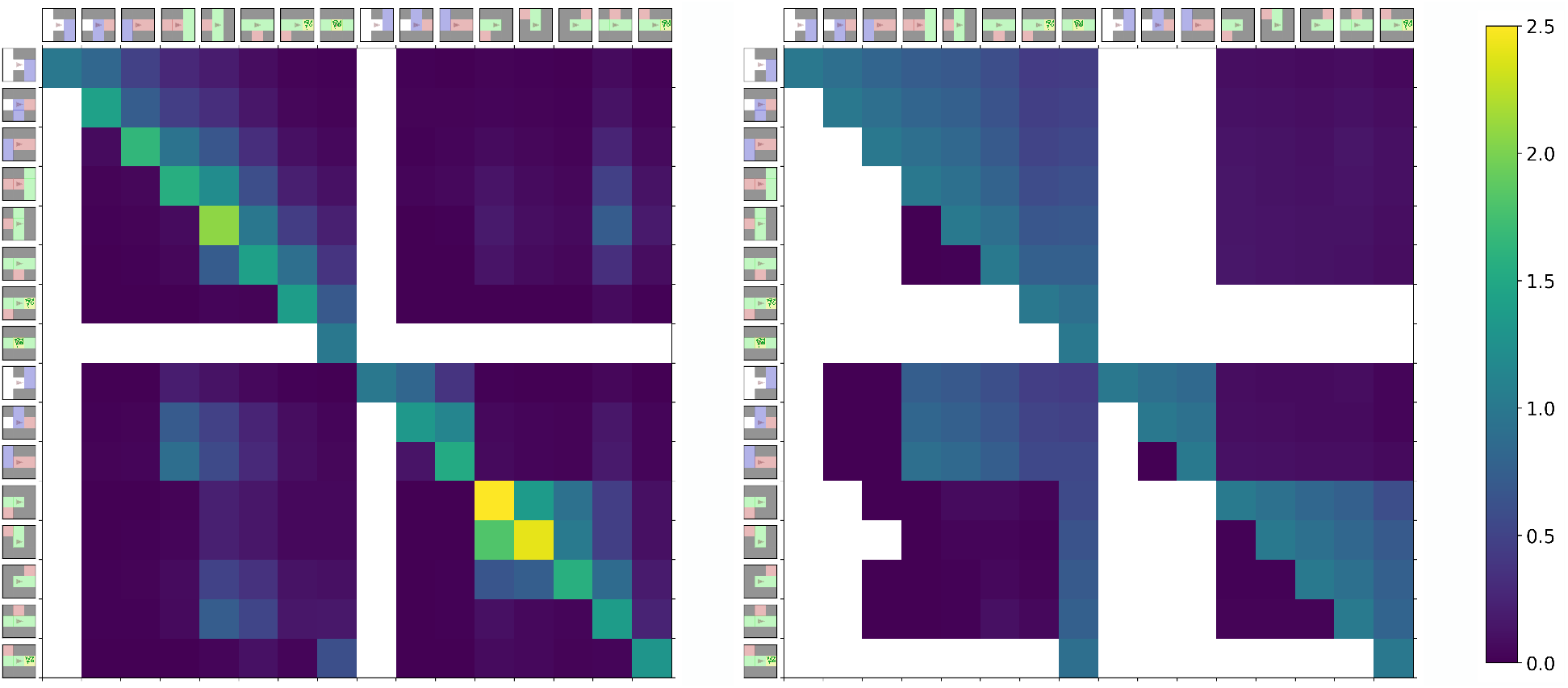
Successor Representations sub-matrices. We show two SR sub-matrices constructed from the state transitions experienced while learning to habituate during the first phase of learning (left matrix 90% of the actions is random, *ε* = 0.9) and final phase (right matrix *ε* = 0). Each cell value corresponds to the future discounted visitation expectancy of the column state given that we are in the row state. Notice that the scale goes up to 2.5 in the left matrix meaning that when getting stuck in one corner we expect to visit more than once those contiguous states. If a cell is filled white, it means that its value was less than 10^−5^ (this is indicative of a never or very rarely experienced transition). Towards the end of the learning (right matrix) the scale collapses to the range 0…1 meaning that we do not expect to visit the same state twice.

After a certain number of learning iterations (episodes), the absolute value of the 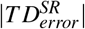 signal, that is |*γ^SR^ − M_t_*(*S_t_*, *S*_*t*+1_)|, will provide the surprise signal that we were looking for (see Section 2.6). When this quantity will be greater than a threshold (we will use 0.5), it will mean we are highly surprised of being in state *S*_*t*+1_. As we show in Figure **12**, the SR *TD*-error signal is suited to provide the surprise signal that we need to recognize and be aware of the anomaly. Well in fact, after an habituation process, if we freeze the learning, the 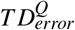 of Q-Learning, which is a reward based surprise signal will also be very high when experiencing the anomaly (as we also plot in Figure **12**). So here a discussion could start about the need of another surprise signal. We think the awareness of the anomaly is a Model Based signal, as we identify the state as being not expected. If we were in the beginning of learning, the need of an alternative SR based signal would be evident. The SR *TD*-error signal could in fact modulate the Dyna-Q model updates, solving the convergence problems experienced under partial-observability. We leave this idea for future investigations. Another interesting question is if both TD errors follow a linear relationship during learning. We answer this question negatively by providing the scatter plot of both errors during learning in Figure **12b**. But we provide an additional result in favor of the two parallel processes by looking at how both TD errors evolve over time in Figure **12a**. The Q-Learning TD-error 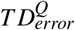 increases at the end of learning because of the need of re-adjusting all the errors associated to the new trajectory when experiencing a rare start into the bottom position. On the other hand, the variability (standard deviation) of the SR TD-error reduces with time, providing a more consistent signal, the reason being we traverse fewer and fewer states as learning progresses.

**Figure 12:**
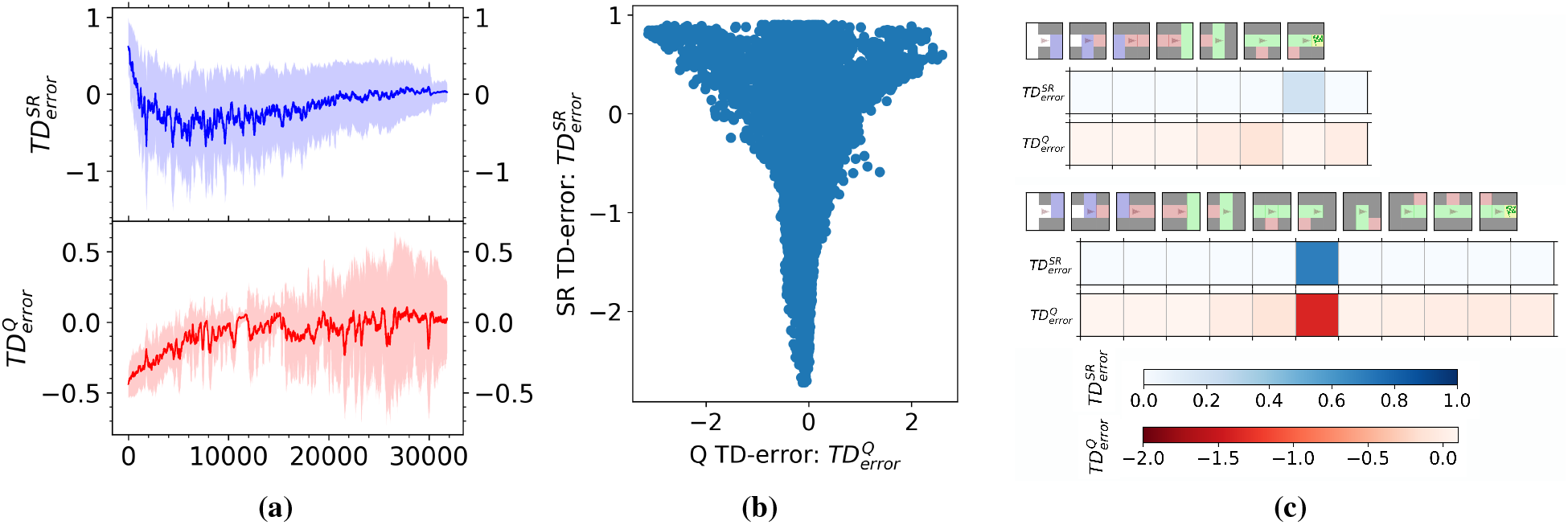
Successor Representation and Q-Learning TD-error comparison. **(a) (b) (c)** Shows two sequences of state visits after learning (from restart position 1 and 2) and their associated SR surprise signals |*γ^SR^ − M*(*S_t_, S*_*t*+1_)|. When taking the uncommon restart position 2 (right elevator side) and experiencing the HER anomaly (bottom trajectory), both error SR and Q-Learning TD-errors are higher and lower respectively.

SRs would provide an explanation of how are we able to detect the anomaly and find ourselves in the wrong state or knowing that we should have arrived to our goal, but how we do solve the anomaly? We will deal with this in next Section 3.9.

### 3.9 Retrospective inference of the cause of the habituation failure

We reduced the HER task to a One-Choice task with partial observability and we have compared it to the task in [27] in Section 3.1. In [27] a retrospective inference mechanism is described and identified in behavioral experiments. When facing a situation under incomplete information, and having the outcome of an expected reward delivery, participants perform a model-based inference by reinforcing a state that was not directly observed (the state to be reinforced is inferred from a Model-Based learned structure of the task). The surprise signal is driven in that case by reward expectancy as reward is non-stationary (usually in all Two-Choice and One-Choice tasks [2]). The 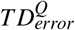 of Q-Learning already carries a reward expectancy signal that is in fact used by experience replay and prioritized sweeping [21] mechanisms.

In previous section we modeled a surprise state visitation signal (based on Successor Representations) that allows to detect the anomaly and be aware of the undesired situation. When the surprise signal is sufficiently high and we found ourselves in a state that was totally unexpected (that is when |*γ − M_t_*(*S_t_, S*_*t*+1_)| > 0.5), we activate a modelbased goal-directed behavior. We hypothesize that a retrospective model-based explanation mechanism (or inference as called in [27]) must be in place to try to explain the source of this habituation failure. We also hypothesize that the source of error is attributed to the last decision made and that corresponds in the HER task to the exit of the elevator). However the question arises: how to distinguish a relevant environment behavioral decision point when in the RL formulation we take an action and decision at every step? This can be reformulated with the question: what RL subset of actions in a state are relevant decision points? Not all RL actions available at a state represent relevant environment behavioral decision points. As explained in Section 3.3, this past state could also be identified using the Value of Perfect Information (VPI) measure (or our customized version described in Section 3.3). The brain raises an alarm signal to the last highest VPI experienced state, although, as we discussed, VPI is not a sample efficient measure.

### 3.10 One shot learning with retrospective alarm model-based switch

The retrospective Model-Based mechanism assigns an alarm signal at the decision point. So the state *S*^*r*=1^ corresponding to the identified source of error causing the anomaly (identified by the SR surprise signal) gets assigned an increased alarm marker that we call *Z*(*S*^*r*=1^), which is increased.

When facing again the same situation (at the next episode), when the agent is at the decision state *S*^*r*=1^, the alarm signal *Z*(*S*^*r*=1^) will be non-zero and will activate a Model-Based increase attentional response (available as an action in the *Observation Size On-demand* environment) which will return a state with radius 2, *S*^*r*=2^). As the learning has been carried out by the so-called onion updates (described in Section 3.7), the Model-Free mechanisms has an optimal response to that increased state size *S*^*r*=2^. Therefore, a one-shoot learning response can be learned that achieves the optimal policy under partial observability: activate the increased observation response at the decision state (when exiting the elevator).

In the following we show the policy that agent is executing in the case of the one shot reaction to the HER anomaly:

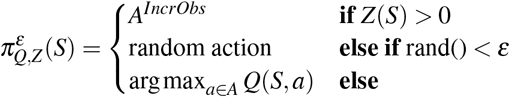

If the alarm signal is non-zero we perform an increased observation demand action to the environment (losing one step, thus obtaining −0.5 reward). If this is not the case we perform a random action with probability *ε* and we exploit the current state-action value function otherwise by “greedifycation” (see Section 2.3). The alarm arbitration mechanism is a Model-Free mechanism that triggers an increased attentional action given a state: *Z*(*S*^*r*=1^) and it is a less demanding memory mechanism which remembers that something bad happened in that state, thus a switch to Model-Based (in this case, increased attention) should be made.

### 3.11 Model-Based landscape

We also summarize in the high level schema of Figure **13** the underlying MF/MB arbitration process that we propose is driving human behavior in the HER task. We also characterize as a sort of final discussion to the paper how Model-Based processes can be interpreted in different forms: sub-goal task division, increased attentional demands for enhanced state information, retrospective inference (as in [27]) for explaining the source of habituation failures (Section 3.9) and reconsidering decisions and forward planning (as in [18]).

**Figure 13:**
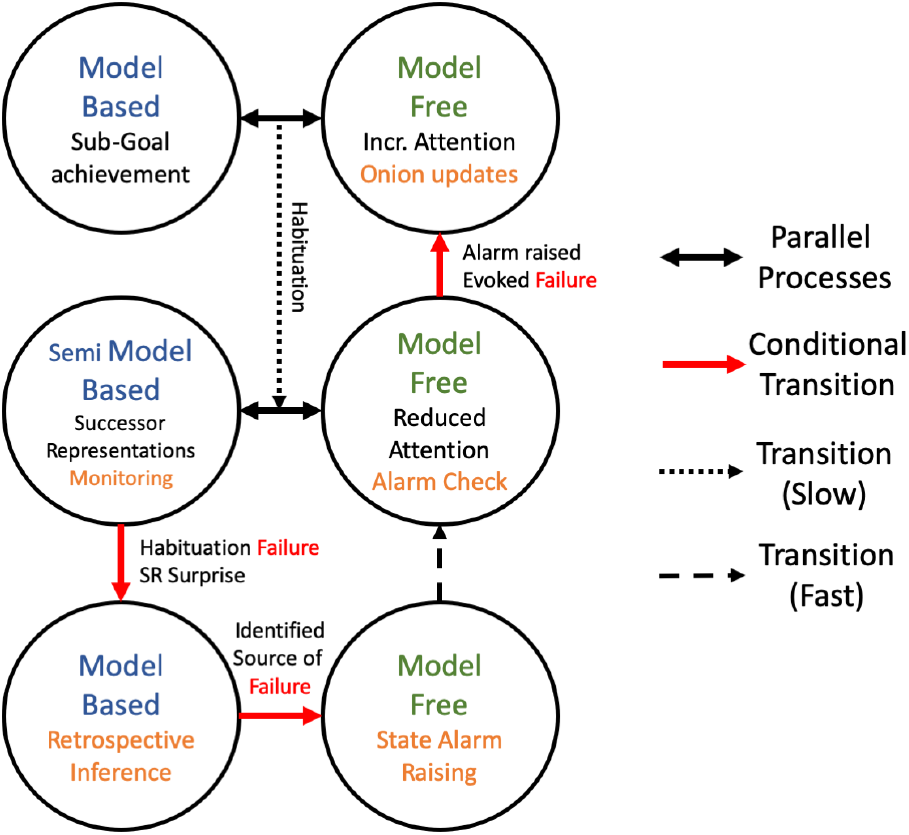
Model Free / Model Based arbitration mechanisms. First row: parallel processes described in Section 3.7. Second row: parallel processes described in Section 3.8. Third row: described in Sections 3.9 and 3.10.

## 4 Conclusions

We have defined and modeled within the Reinforcement Learning framework, the so-called *Hotel Elevator’s Row* (HER for short). Briefly, the HER anomaly is described as finding yourself in the wrong corridor, when exiting the elevator in hotels with two elevator’ sides. HER is a partial observable task that captures a recognizable human behavioral conflict between habitual and goal-directed behavior in which the habituation process may fail repeatedly due to incorrect fast decision-making under incomplete information. There is an efficiency-flexibility trade off between Habituation (Model-Free), Goal-Directed (Model Based) and in-between approaches like Successor Representations (SRs) [15]; being habituation more efficient and Goal-Directed behavior more adaptable to environmental modifications and changing reward contingencies [28]. We have shown by means of the HER task, that this trade off gets more complex under partial observability, and thus the need to identify and model new arbitration mechanisms between these systems as we have done. Lets mention also that it is not clear what are the limits of habituation, as even model-based behavior can become automatic [2, 11].

We have proposed four different variants of the HER task that include different aspects, important to match its human behavioral counterpart. (1) The usual environment with global, complete information states (used in nearly all studies ([21, 28] for instance), (2) An environment with an agent that has a reduced observability around him, having access only to incomplete information / partial observable states (as introduced in [27] and also in [33] with another type of partial observability, noisy perception), (3) An agent being able to remember a window of past observations, thus having memory capabilities (memory seems to be the only described remedy for dealing with partial observability [23]), (4) A complete/incomplete observation size on-demand by adding an additional action in the RL formulation which loses one time step but delivers a higher resolution observation (this variable resolution observation is new to our knowledge in RL formulations, but previously introduced in the planning community).

When considering the usual complete information or memory environments (the just mentioned n°1 and n°3), the HER task is a prototypical RL task in which the anomaly is not reproduced and Model-Based approaches like Prioritized Sweeping DynaQ outperforms Model-Free Q-Learning (DynaQ is more sample efficient, with less interactions with the environment it finds an optimal policy).

As we have shown, the picture changes completely when introducing partial observability (in the usual way with a limited perception radius around the agent as we do in environment n°2 or with an observation size on-demand (including an additional RL action that wastes one step but delivers a larger observation as in environment n°4). Then the HER anomaly (one of the learned trajectories always includes an undesired detour due to partial observability) is reproduced with all RL algorithms (MF Q-Learning, MB DynaQ, in between SR-Learning and Reinforce Policy Gradient). It happens that state of the art Model-Based methods like Prioritized Sweeping DynaQ (used in [21] to explain underlying physiological processes) have a worst performance than Model-Free Q-Learning in this setting and fails to converge (thus the plural failures in the title of the paper).

We are permanently exposed to partial observability (incomplete information) and the brain is obliged to detect ambiguous and possible future undesired situations and take decisions including the right minimal contextual information at the lowest possible cost. At a computational level, to benchmark RL and Deep RL algorithms environments and scenarios like Animal-AI Olympics^5^, Obstacle Tower Challenge^6^, and all first person view games are intrinsically partial observable as they deliver a first person view of the world state (our egocentric agent’s observation is analogous to first person view) and many other games, like the Atari game suite used in [22], suffer from certain degrees of partial observability. Moreover, when using Deep Neural Networks to approximate Reinforcement Learning elements (Value Functions, Policies, etc.), the network creates a non-controlled partial observability that can lead to convergence learning problems and undesired situations similar to the one expressed minimally by the HER anomaly.

We have gone further in proposing various hypothesis on the underlying Habituation / Goal-Directed arbitration mechanisms (going beyond the MF/MB dichotomy [4]), that could be at play when behaviorally identifying the HER anomaly and trying to avoid it.

For the mechanisms that enable the behavioral identification and awareness of the anomaly, we propose and present simulation evidences for MF and MB parallel processes (as in [30], in our case, learning to habituate while being monitored by semi Model-Based state expectancy visitations). A Model-Based Successor Representations mechanism that acts as a mismatch detector could operate at several dimensions, delivering alternate surprise signals: states expected to be visited and also in the correct timing. Once the habituation failure (the HER anomaly) is identified, the brain triggers a Goal-Directed mechanism to find the sources of that highly surprising situation. We call this process retrospective inference (as in [27]) and we have hypothesized that the brain identifies and reconsiders the last state in which we made a relevant decision. We further hypothesize that an alarm signal (Model-Free mechanism) is reinforced in that source state to mark it as a cause of habituation failures and switch to a Goal-Directed MB behavior when encountered again in the future. Summarizing, we propose that habituation operates together with a Sucessor Representation monitoring mechanism that provides a surprise signal that activates Goal-Directed behaviors. In addition, a model-based retrospective inference, assigns an alarm signal to crucial decision making states in which habituation (that failed in the past) is inhibited and goal-direct behavior activated.

Many people can recognize to have experienced the HER anomaly, but systematic behavioral experiments could also be designed to provide more evidences of the different proposed arbitration mechanisms. One can think of a realistic Virtual Reality task that would mimic the HER environment or even to make use of the simplistic observations and analogous task described here. Exposing people to increased cognitive load during the elevator’s traversal would be necessary to avoid people using the remember/memory strategy (which we proven solves the anomaly and avoids partial observability). Analyzing reaction times during action execution at each state, one could match the moment of awareness of the anomaly and also try to find behavioral correlates of the retrospective inference mechanism that, we described, assigns an alarm signal to the state identified as the source of the error. Also this experiments could really discern if humans solve this task in few shot learning episodes after experiencing the anomaly. This behavioral experiments could provides clear directives to look for the existence of such processes in the brain and to understand their neuro-physiological basis as we did in [12] with the human study with intracranial LFP analysis during the realization of a Virtual Reality task.

The identified, described and modeled mechanisms have as well clear applications at a computational side as algorithms could be designed with these principles: for example utilizing several sources of surprise signals in addition to the usual reward based temporal difference error other model-based state visitation and timely expectancy, can be added.

As other research on MF/MB arbitration shows, the identification of these new mechanisms could shine light into new treatments for addiction (as the proposed Model-Based monitoring systems could be trained to reduce it), compulsive behaviors like compulsive checking (again fostering the parallelisation of Model-Free Model-Based systems) and understand better car accidents caused by attention deficits due to habituation (providing hints on how to desautomatize habits, by training Model-Based processes as well).

## Acknowledgments

Research supported by INSOCO-DPI2016-80116-P. This paper, benefited from the fruitful discussions with Clément Moulin-Frier, Vicenç Gómez and Daniel Pacheco.

## Footnotes

3 As mentioned in [32] Section 14.6 page 368

4 See for example Section 14.6 of [32] page 367

5 http://animalaiolympics.com/AAI/

6 https://github.com/Unity-Technologies/obstacle-tower-env

